# Growth of the Radiotrophic Fungus *Cladosporium sphaerospermum* aboard the International Space Station and Effects of Ionizing Radiation

**DOI:** 10.1101/2020.07.16.205534

**Authors:** Graham K. Shunk, Xavier R. Gomez, Christoph Kern, Nils J. H. Averesch

**Affiliations:** Higher Orbits “Go For Launch!” Program, Leesburg, VA, United States; Physics Department, North Carolina School of Science and Mathematics, Durham, NC, United States; Department of Systems Engineering, University of North Carolina at Charlotte, NC, United States; Department of Statistics, Ludwig Maximilian University of Munich, Munich, Germany; School of Social Sciences, University of Mannheim, Mannheim, Germany; Department of Civil and Environmental Engineering, Stanford University, Stanford, CA, United States; Center for the Utilization of Biological Engineering in Space, Berkeley, CA, United States

**Keywords:** Space, Radiation-Shield, Fungus, *Cladosporium sphaerospermum*, Radiotrophic, ISRU

## Abstract

The greatest hazard for humans on deep-space exploration missions is radiation. To protect astronauts venturing out beyond Earth’s protective magnetosphere, advanced passive radiation protection is highly sought after. In search of innovative radiation-shields, biotechnology appeals with suitability for *in-situ* resource utilization (ISRU), self-regeneration, and adaptability.

Certain fungi thrive in high-radiation environments on Earth, such as the contamination radius of the Chernobyl Nuclear Power Plant. Analogous to photosynthesis, these organisms appear to perform radiosynthesis, utilizing ionizing radiation to generate chemical energy. It has been postulated that the absorption of radiation is attributable to the pigment melanin. It is further hypothesized that this phenomenon translates to radiation-shielding properties.

Here, growth of *Cladosporium sphaerospermum* and its capability to attenuate ionizing radiation, was studied aboard the International Space Station (ISS) over a period of 26 days, as an analog to habitation on the surface of Mars. At full maturity, radiation beneath a ≈ 1.7 mm thick lawn of the dematiaceous radiotrophic fungus was approx. 0.84% lower as compared to the negative-control. In addition, a growth advantage in Space of ∼ 21% was observed, substantiating the thesis that the fungus’ radiotropism is extendable to Space radiation.

## Introduction

### Background

With concrete efforts to return humans to the Moon by 2024 under the Artemis program and establish a permanent foothold on the next rock from Earth by 2028, humankind reaches for Mars as the next big leap in Space exploration [1]. In preparation for prolonged human exploration missions venturing beyond Earth-orbit and deeper into Space, the required capabilities significantly increase [2]. While advanced transportation solutions led by the public and private sectors alike (SLS/Orion, Starship, New Glenn) are pivotal and have already reached high technological readiness, life support systems as well as crew health and performance capabilities are equally essential. Therefore, any mission scenario (cf. ‘Design Reference Architecture 5.0’ [3] or ‘Mars Base Camp’ [4]) must include innovative solutions that can meet the needs and address the hazards of prolonged residence on celestial surfaces.

The foremost threat to the short- and long-term health of astronauts on long-duration deep-Space missions is radiation [5, 6]. Over the period of one year, the average person on Earth is dosed with about 6.2 mSv [7, 8], while the average astronaut on the International Space Station (ISS) is exposed to an equivalent of approximately 144 mSv [9]; one year into a three-year mission to Mars, an astronaut would already have accumulated some 400 mSv, primarily from Galactic Cosmic Radiation (GCR) [10]. While the particular health effects of radiation exposure on interplanetary travel have not been fully assessed [11], it is clear that adequate protection against Space radiation is crucial for missions beyond Earth-orbit. Both active and passive radiation-shields, the latter investigating inorganic as well as organic materials, have been and are extensively studied [12, 13], but solutions are more than any other factor of Space travel restricted by up-mass limitations [13].

Hence, *In-Situ* Resource Utilization (ISRU) will play an integral role to provide the required capabilities, as well as to break the supply chain from Earth and establish sustainable methods for Space exploration, because once underway there virtually is no mission-abort scenario [14]. For ISRU, biotechnology holds some of the most promising approaches [15-19], posing useful for providing nutrition [20], producing raw-materials [21] and consumables [22], and potentially even “growing” radiation shielding [23].

Among all domains of life exist extremophiles that live and persist in highly radioactive environments, including bacteria, fungi, as well as higher organisms such as insects [24, 25]. Certain fungi appear to be able to utilize ionizing radiation through a process termed radiosynthesis [26], which is perceived analogous to how photosynthetic organisms gain chemical energy from electromagnetic radiation of the visible spectrum [27, 28]. Large amounts of melanin in the cell walls of these fungi protect the cells from radiation damage while presumably also mediating electron-transfer, thus allowing for a net energy gain [29, 30]. Melanized fungi have been found to thrive in highly radioactive environments such as the cooling ponds of the Chernobyl Nuclear Power Plant, where radiation levels are three to five orders of magnitude above normal background levels [31]. Additionally, they populate the interiors of spacecraft in low Earth orbit (LEO), where exposure to ionizing radiation is also intensified [28]. Black molds and their spores have been found to remain viable even after exposure to an equivalent radiation dose of several months’ worth of Space radiation [32]. How these organisms protect themselves from radiation damage has been the subject of intense study and specifically melanin has been explored as biotechnological means for radiation shielding [33, 34].

Here, we explore the physiological response of a melanized fungus to the environment of the ISS and assess its potential to serve as bioregenerative part of a multi-faceted passive radiation-shield.

### Concept

The preference of *Cladosporium sphaerospermum* (a dematiaceous fungus, cf. supplementary information 1, section A) for environments with extreme radiation-levels on Earth is well documented [27, 28, 35]. Consequently, it has been hypothesized that similar proliferation occurs in response to the high radiation environment off-Earth and that such melanized fungi can be utilized for radioprotection in Space [23, 34]. The objective of this experiment was to conduct a proof-of-principle study on a single payload, utilizing basic flight-hardware for an autonomous experiment in the unique radiation environment of the ISS. This offers the opportunity to test the fungus’ (growth) response in Space, and the capacity of the formed biomass to attenuate ionizing radiation. In an additional analysis based on the concept of linear attenuation (Lambert’s law) [36], the specific contribution of melanin to provide adequate shielding against ionizing radiation was assessed.

## Materials & Methods

### Experimental Setup

Space Tango (Space Tango, Inc., Lexington, KY, US) was contracted for experimental design and construction (terrestrial logistics and on-orbit operations) [37]. The initial concept for the experimental design was adapted by Space Tango for assembly of the flight-hardware and implementation aboard the ISS within TangoLab™ facilities. The flight-hardware was housed in a 4”×4”×8” double unit standard-size CubeLab™ hardware module and consisted of the following main components: two Raspberry Pi 3 Model B+ (Raspberry Pi Foundation, Caldecote, Cambs., UK) single-board computers, EP-0104 DockerPi PowerBoard (Adafruit Industries, New York, NY, US), [PocketGeiger Type5](http://www.radiation-watch.org/2011/05/professional.html) (Radiation Watch, Miyagi, JP) with the PIN photodiode X100-7 SMD (First Sensor AG, Berlin, DE), Raspberry Pi Camera v2 (Raspberry Pi Foundation, Caldecote, Cambridgeshire, UK) and light source (0.8 W LED-strip) for imaging, DHT22 integrated environmental sensor suite (Aosong Electronics Co. Ltd, Huangpu District, Guangzhou, CN) for temperature and humidity readings, a real-time WatchDog™ timer (Brentek International Inc., York, PA, US), and [D6F-P0010A1](https://omronfs.omron.com/en_US/ecb/products/pdf/en-d6f_p.pdf) (Omron Electronics LLC, Hoffman Estates, IL, US) electronic flow-measurement system. One Raspberry Pi (“auxiliary-computer”) running Raspbian v10.18 was dedicated to photography, lighting, temperature, humidity, and electronic flow measurement (EFM) readings, while the second Raspberry Pi (“flight-computer”) controlled radiation measurements, stored in a probed Logger Memobox (Fluke Corporation, Everett, WA, US). The assembled flight-hardware was calibrated and vetted before flight; in particular consistency of the two radiation sensors was confirmed so that no deviation in recorded counts existed between them.

*Cladosporium sphaerospermum* (ATCC® 11289™) was obtained from Microbiologics (St. Cloud, Minnesota, US), catalog no. 01254P. Growth medium was potato dextrose agar “PDA” (Carolina Biological, Burlington, NC, US) obtained as “Prepared Media Bottle” (approx. composed of 15 g/L agar, 20 g/L glucose, and 4 g/L starch). A total of 20 mL PDA (dyed with orange 1) was used to fill the two compartments of a split Petri dish (100×15 mm). The agar plate was sealed with ‘Parafilm M’ post-inoculation. With a total height of the Petri dish of 15 mm and a 75 cm^2^ surface area, the thickness of the PDA was ∼ 13.33 mm, leaving an approx. 1.67 mm gap for the fungal growth layer. Cellular mass density of fungal biomass (1.1 g/cm^3^) [38] was assumed to remain unaffected by microgravity. To minimize latent growth while in transit, the setup was fully assembled before the inoculation of the medium. A block-chart of the experimental flight-hardware setup is given in figure 1, further details are provided in supplementary information 1, section B.

**Figure 1:**
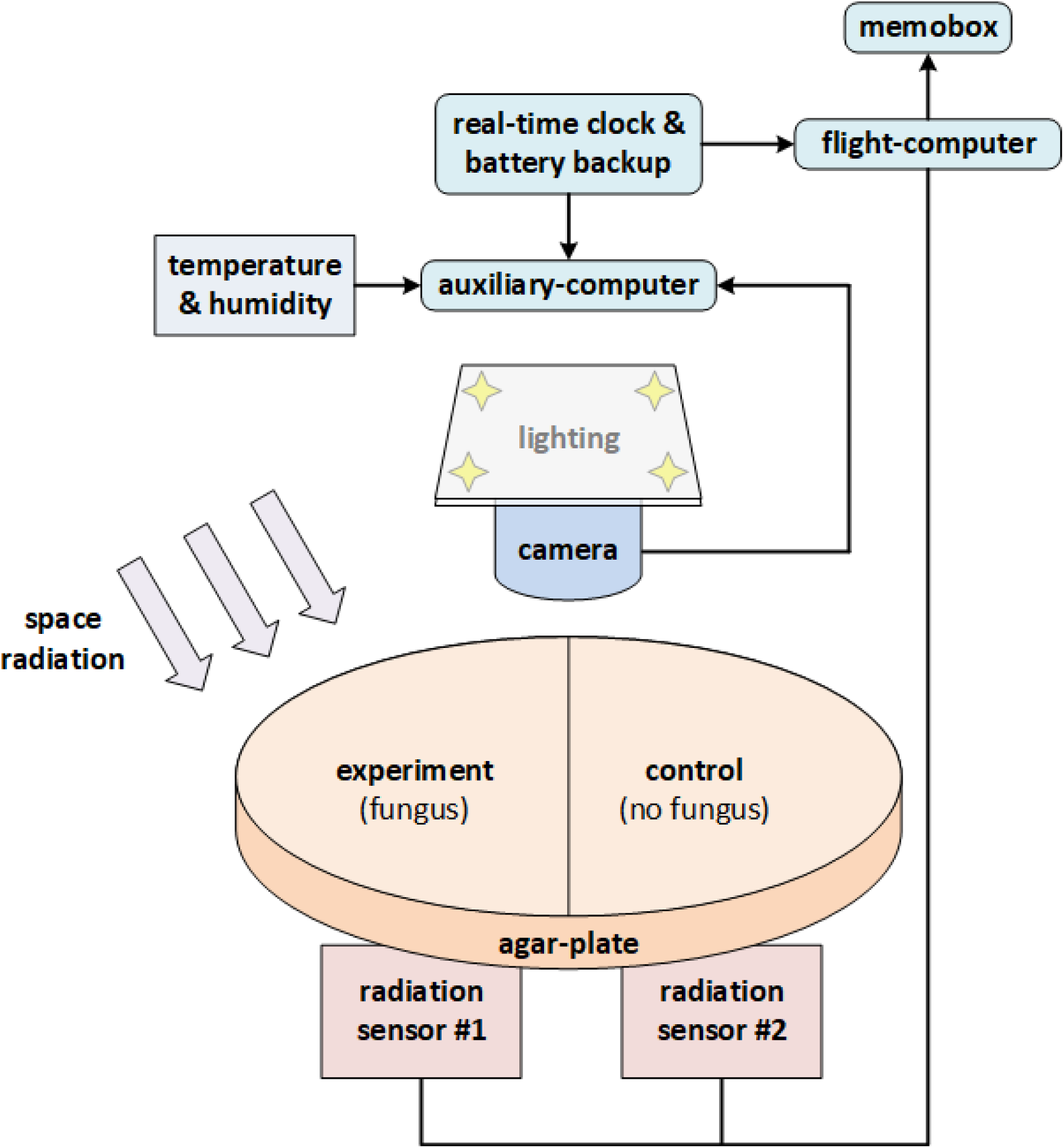
Block-chart of the experimental flight-hardware setup. The single (split) Petri dish accommodated both the experiment (agar + fungus), as well as the negative-control (agar only). The two (square) radiation sensors were situated in parallel directly beneath the Petri dish (one for each side). Note that “shielding” is one-sided only (for simplicity of experimental setup and to keep mass low).

### Vetting for Cold-Stow

The response of *Cladosporium sphaerospermum* to cold storage was determined in a preliminary experiment. A total of six Petri dishes containing PDA were inoculated with the fungus, five were stored at 4°C with one kept at room temperature (RT) as control. Plates were removed sequentially after 1, 5, 10, 15 and 20 days. Fungal growth on each plate was monitored at RT and compared to the control.

### On-Orbit Implementation

The equipment was packaged with the inoculated Petri dish before flight, accommodated in a [2U CubeLab™](https://spacetango.com/cubelab/) (Sealed) [39], between Dec 2018 and Jan 2019. Lead-time before the launch was two days in cold-stow. Transition time to LEO was three days (in total approx. one week to “power-on” after inoculation), transported to the ISS (U.S. Destiny Laboratory) in cold storage (4°C) on SpaceX mission CRS-16. On orbit, the experiment was oriented such that the Petri dish (and radiation sensors) faced away from Earth. Pictures of the Petri dish were taken in 30-minute intervals for 576 hours, resulting in 1,139 images. Temperature, humidity and radiation were measured every ∼ 1.5 minutes throughout the 622-hour run-time of the experiment. The three temperature sensors recorded 27,772 measurements each. Radiation was measured incrementally; 23,607 radiation and noise-counts were recorded by each radiation sensor. The stand-alone lab was active aboard the ISS for ∼ 26 days with data downlinked ten times, on average every two to three days, before power was cut and the experiment awaited return to Earth in June 2019.

### Ground-Controls

In addition to the preflight growth test and integrated on-orbit negative-control, Earth-based growth experiments were performed post-flight, replicating the conditions of the flight-experiment without radiation. The same methods and techniques were utilized when preparing the cultures on solid medium. As means of a ground-control, a time-dependent temperature profile analogous to the on-orbit experiment was replicated and graphical data was collected in the same intervals, to record the growth behavior.

### Evaluation of Growth

Photographs of the cultures were processed with MATLAB [40] (The MathWorks Inc., Natick, MA, US) to derive the average brightness values for congruent subsections of each image (cf. supplementary information 1, section C), as proxy for biomass formation. These average brightness values were then normalized to render the relative optical density (OD) ranging from zero to one (cf. supplementary information 2) over time. Exponential regression models (implemented in R, cf. supplementary information 4) allowed growth rates “*k*” to be determined. Based on these, a relative difference (in percent) between *k* _*exp*_ for the on-orbit experiment and the ground-control, *k*_*ctrl*_, was estimated.

### Evaluation of On-Orbit Radiation

Cumulative radiation counts, derived from the incremental data recorded by the radiation sensors of control- and fungus-side, were plotted over the runtime of the on-orbit experiment. The relative difference in radiation attenuation (in percent) between negative-control and fungus-side was determined based on the slope of linear regressions in different phases, which were defined based on the relative OD. Phase 1, the initial phase, was defined for relative OD measures below 50% of the maximum, corresponding to the first 19 hours. Phase 2, the growth phase, was defined for relative OD measures between 5% and 95% of the maximum, correspondingly starting 5 hours after the start (t_0_) and ending 46 hours after t_0_. Phase 3, the stationary phase, comprised data from 200 to 622 hours after t_0_, corresponding to relative OD measures greater than 99% of the total maximum.

### Radiation Attenuation Analysis

Applying Lambert’s law, the concept of linear attenuation served to complement the experimental part, focusing on ionizing electromagnetic radiation. The analysis determined the attenuation capacity of the fungal biomass within the absorption spectrum (low keV range [41]) of the employed radiation sensors (cf. supplementary information 1, section B).

To interpret the results in light of a more relevant (real-world / Space radiation environment) scenario, experimental findings were used to inform a theoretical analysis. This relied on the extensive resources that exist for attenuation coefficients of different substances over a range of photon energies by means of the NIST X-ray Attenuation Databases [42]. Particularly, mass-attenuation coefficients (MACs) for the respective compounds and mixtures (cf. supplementary information 3) were derived from [NIST-XCOM](https://physics.nist.gov/PhysRefData/Xcom/html/xcom1.html) for total coherent scattering. The non-linear influence of secondary radiation was respected by means of buildup-factors [43] and equivalent kinetic energies of subatomic and elementary particles (in eV) were used in the calculations. The correlations and assumptions, as well as the specific workflow used in the attenuation analyses are described in detail in supplementary information 1, section D & E.

### Validation of Statistical Robustness

Logistic growth-curve models were fitted to relative OD measures from the ground-controls and on-orbit experiment, to test for differences in the slopes of the curves and hence growth behavior. Differences in radiation counts over time between the experiment and the negative-control of the on-orbit experiment were modeled with a set of robust regressions. Data preparation and model specification are outlined in supplementary information 4, section A.

## Results & Discussion

### Pre-Flight -Cold-Stow Growth-Test

The ‘Cold-Stow’ experiment showed that for all refrigerated sample-plates there was insignificant fungal growth immediately upon removal from incubation at 4°C (i.e., no fungal growth prior to t_0_) for all trialed timeframes. Furthermore, all samples exhibited similar growth (data not shown) once at ambient temperature, regardless of the time spent in cold storage.

### Microbial Growth Advantage On-Orbit

From the point the hardware was powered on aboard the ISS, the temperature rose sharply from the initial 22°C, reaching 30°C within 24 hours and stabilized after 48 hours around 31.5±2.4°C for the rest of the experiment (cf. supplementary information 2).

Many dimorphic fungi are characterized by slow growth and require up to 14 days for significant biomass formation to occur at an optimum temperature around 25°C [44]. In the on-orbit lab, *C. sphaerospermum* reached maximum growth and full coverage of the PDA already after two days, as discernible from the photographical data (cf. supplementary information 1, section B) as well as the derived growth-curve, shown in figure 2. Comparison with the ground-controls indicates that the fungus may have experienced faster-than-normal growth aboard the ISS, as per the modeled growth-curves (supplementary information 4). Specifically, the growth rate in the on-orbit experiment was on average 1.21± 0.37-times higher (based on *k*_*ground_1*_ = 0.241 h^-1^, *k*_*ground_2*_ = 0.231 h^-1^, *k*_*ground_3*_ = 0.196 h^-1^ and *k*_*flight*_ = 0.299 h^-1^, for the ground-controls and on-orbit experiment, respectively).

**Figure 2:**
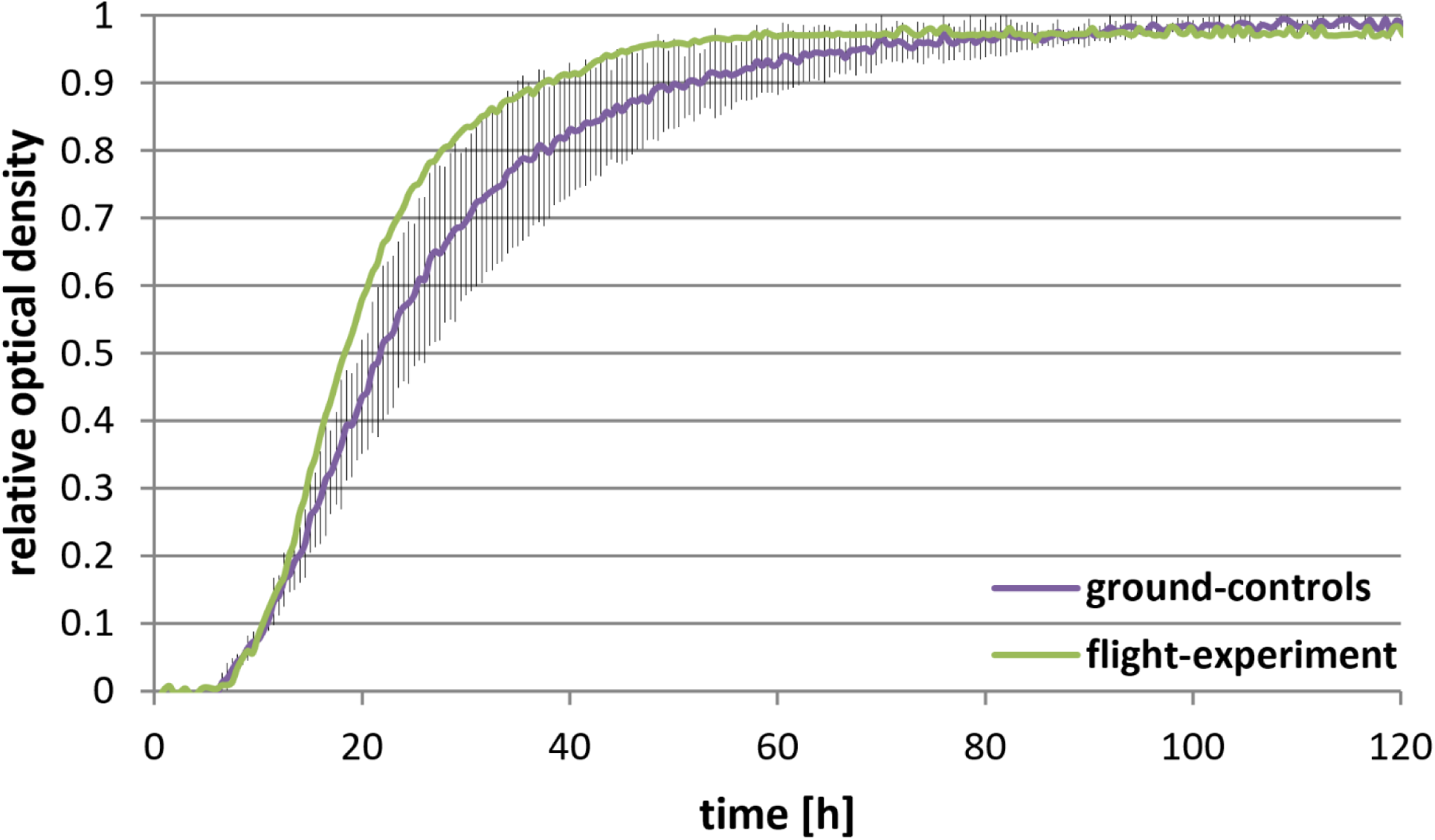
Growth-curves (initial 120 hours shown) for *C. sphaerospermum* cultivated on solid medium (PDA), depicted by means of connected data-points of relative optical densities. While the on-orbit experiment (green) was a single run, the ground-control growth-experiment (purple) was conducted three times (two replicates in each of the three runs). The error bars show the standard deviation between the three separate runs. t_0_ was normalized based on the onset of exponential growth of the flight experiment.

While not immediately numerically tangible, it is worth mentioning that in preliminary ground-control experiments often poor growth was observed at 30°C, as compared to RT (data not shown, not used to establish the ground-control growth-curves), with only sporadic coverage of the PDA with fungal colonies (cf. supplementary information 1, section B). Only experiments where full coverage with *C. sphaerospermum* was achieved were selected as representative ground-controls, even then, inconsistency in growth existed, as is also reflected by the large deviation between the three replicates (error bars depicted in figure 2, cf. also raw-data and additional plots in supplementary information 2). The observation that on Earth higher-than-optimum incubation temperatures rapidly hindered growth is in accordance with literature [45], and strengthens the conclusion that the Space environment benefited the fungus’ overall physiology, where growth was strong and consistent.

The growth advantage in Space may be attributable to the utilization of ionizing radiation of the Space environment by the fungus as a metabolic support function, analogous to Earth-scenarios: when subject to high levels of γ-radiation (500-times stronger than normal) metabolism of melanized fungi appears to be significantly increased, with indication of enhanced growth [29]. Measured by the dose equivalent (144 mSv/a for the ISS and 2.4 to 6.2 mSv/a for Earth [7-9]), the radiation on the ISS is about 20-to 60-times stronger than the average background on Earth, however, about 80% of this is attributable to GCR, which is mostly composed of particle radiation. Hence the fraction of (γ-) rays utilizable by the fungus may not be equivalently significant. In addition, GCR is vastly more damaging [46], which could partially negate the beneficial effect. It is plausible that in addition to direct radiotropism the free-radical scavenging properties of the antioxidant melanin indirectly protected the cells against damage done by radiation, minimizing the detrimental effects inherent to the Space environment [47, 48]. The role of microgravity and impact on growth, whether beneficial or detrimental, cannot be assessed in this context.

### Radiation Measurements in the Space Environment

Due to the nature of the employed radiation sensors (PIN photodiode), dosimetric data was not obtained and no precise statement can be made regarding the absolute radiation levels (e.g., absorbed dose) that the experiment was exposed to. Nevertheless, the periodic fluctuations of the recorded radiation events correlated with dosimetric data from the [U.S. Destiny Laboratory](https://www.nasa.gov/mission_pages/station/structure/elements/us-destiny-laboratory/), specifically those attributed to the transition of the spacecraft through the South Atlantic Anomaly (cf. flightpath of the ISS [49]), as per the trends of the total daily counts (cf. supplementary information 2). Natural phenomena rather than false measurements being the explanation for these incidents is supported by the observation that the spikes of the incremental data coincide for both sensors throughout the entire experiment i.e., high radiation events were picked up by both sensors alike. Consequently, this underscores consistency of the measurements.

### Attenuation of Radiation On-Orbit

Independent of the absolute radiation levels, only the relative difference in ionizing events recorded beneath each side of the Petri dish was significant for the experiment: compared to the negative-control, less radiation counts were recorded directly beneath the side of the Petri dish inoculated with *C. sphaerospermum*, over the runtime of the experiment, as shown in figure 3. To rule out base-deviation of the radiation sensors, a correlation of the radiation counts with biomass formation and change with fungal growth was validated: a significant increase in optical density was only observed after the first day, while no further gain in optical density was observed later than two days into the experiment. Nevertheless, the mark of 200 hours onward was chosen to determine the attenuation capacity to allow for a week of maturation before fungal growth was considered complete and metabolism stationary.

**Figure 3:**
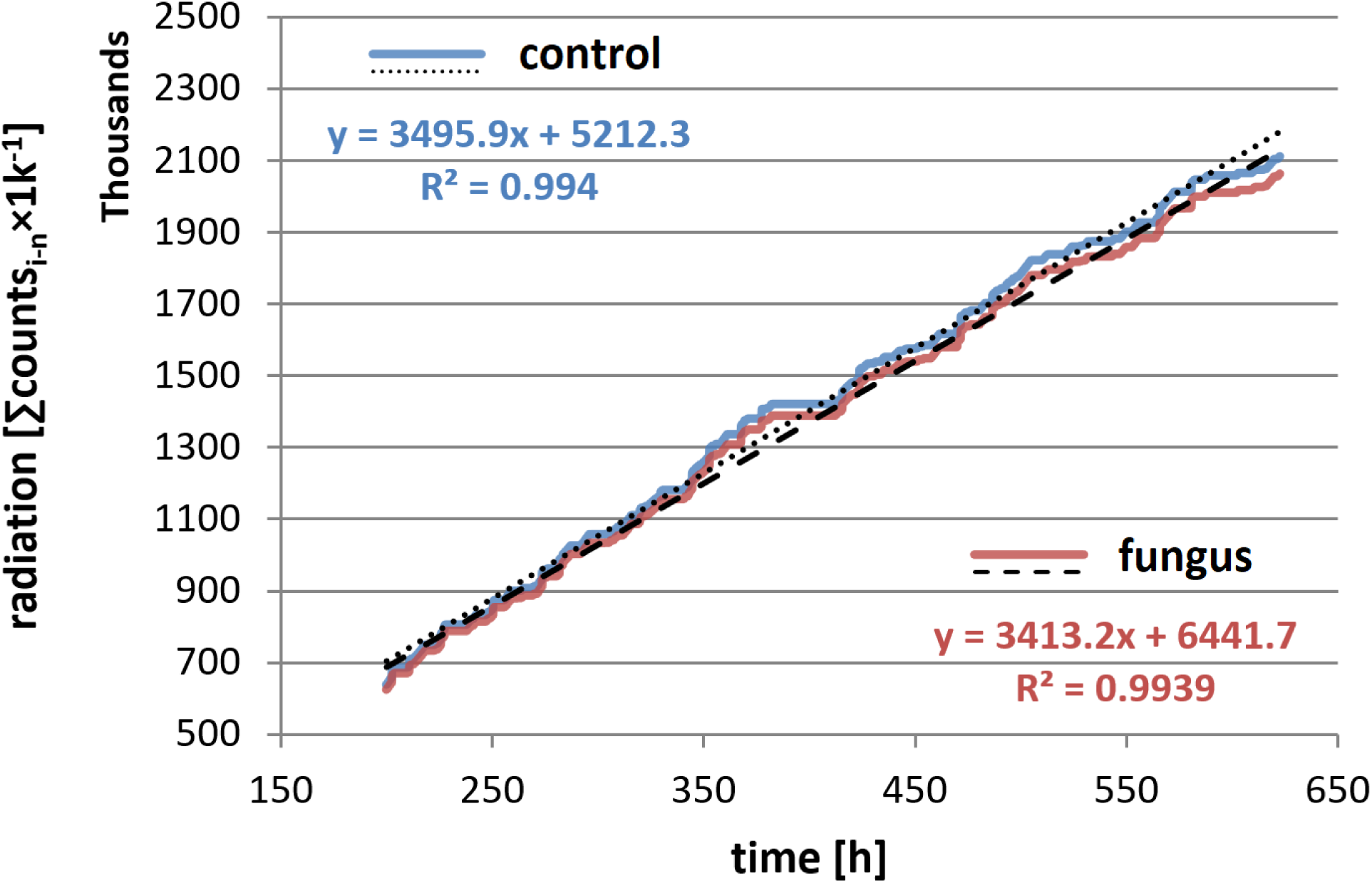
Cumulative radiation counts of the control- (blue) and the fungus-side of the on-orbit experiment (red) over time. (final ⅔ of the experiment, 200 to 622 hours). A significant difference in the slope is evident in section, corresponding to an attenuation of the transmitted radiation. For the sake of legitimacy, the cumulative count of ionizing events (radiation) was scaled-down three orders of magnitude (×1000^-1^).

A greater difference in radiation counts was observable in the late stage of the experiment with full fungal growth as compared to the initial hours when minimal biomass existed: while over the initial 19 hours of the experimental trial radiation counts beneath the fungal-growth side of the Petri dish were approx. 1.53% lower, as compared to the negative-control, the difference during the final 422 hours was on average 2.37% (as per the difference of the linear trendlines’ slopes given in figure 3). This apparent relationship between the amount of fungal biomass (and putatively the melanin content thereof) and change in recorded radiation events could indicate attenuation of ionizing radiation. With a baseline difference between the two sensors of 1.53% (considering that an initial difference between the sensors may have existed or other parts of the hardware constantly shielded a part of the radiation), the observed radiation attenuating capacity can be stated as 0.84%. Since only one side of the radiation sensors was protected by the fungus, it is postulated that also only half of the radiation was blocked. Therefore, it was extrapolated that the fungal biomass may reduce total radiation levels (of the measured spectrum) by up to 1.68%. Considering the thin layer, this may indicate a profound ability of *C. sphaerospermum* to shield against Space radiation of the measured spectrum. Another possible explanation is that conversion of the nutrients contained in the medium into biomass and metabolic end-products and exchange of those with the environment (e.g. consumption of oxygen and production of carbon dioxide) altered the elemental composition of the combined substance in the Petri dish, which resulted in a change of the radiation shielding properties thereof. The full dataset and additional plots of the complete radiation data over the whole course of the experiment can be found in supplementary information 2.

### Statistical Significance and Robustness

Statistical robustness tests were performed with (A) fungal growth data and (B) radiation attenuation data (as provided in supplementary information 2), with the goal to validate significance of the drawn conclusions. Specifically, for (A) it was validated that the slope of the growth-curve based on the flight-experiment data is steeper than the slope of the growth-curves based on the ground-control data and that this difference is statistically significant. For (B) it was validated that there is no significant difference in the average level of radiation measured beneath the negative-control and the fungus/experimental condition in the initial stage of the on-orbit experiment (prior to growth), and the average level of radiation measured beneath the negative-control is higher than beneath the fungus/experimental condition in the late stage of the on-orbit experiment (full growth). This supported the hypothesis that the difference in radiation levels beneath the negative-control and the fungus/experimental may be attributable to the presence of fungal biomass. A detailed description of the methods as well as specific results and interpretation of these can be found in supplementary information 4.

### Radiation Attenuation in Light of ISRU on Mars

The relative difference between the radiation attenuation in the late and early stage of the experiment was used to estimate the linear attenuation coefficient (LAC) of *C. sphaerospermum*, µ_*Fungus*_. The LAC provides a measure for the fungus’ capacity to shield against ionizing radiation and further allowed the estimation of the biomass’ melanin content. Based on this, the required amount (i.e. area density) of melanized fungal biomass that could negate a certain radiation dose equivalent was approximated, to put the shielding potential into perspective. Relevant calculations can be found in supplementary information 1, section D & E.

The LAC of *C. sphaerospermum* at 10 keV was determined to be µ_*Fungus*_= 5.2±4.2 cm^-1^, such that with a density of *ρ*_*m*_ ≈ 1.1 g/cm^3^ for wet fungal biomass [38] the MAC for *C. sphaerospermum* was derived as µ_*Fungus*_*/ρ*_*m*_= 9.51 cm^2^/g. The experimental attenuation coefficient is, however, only valid for the specific γ-energy absorption range of the employed radiation sensors (∼ 10 keV, cf. supplementary information 1, section B). An approximate LAC for melanized biomass at any energy can be determined if the composition of fungal biomass, in particular the melanin content, is known. Here, it was estimated that about 7.2% [w_melanin_/w_CWW_] of the accumulated *C. sphaerospermum* biomass were composed of melanin (cf. supplementary information 1, section D for derivation).^1[50-52]^ The high melanin content of *C. sphaerospermum* is potentially a metabolic response to the strong radiation environment of the ISS, as is consistent with observations of melanized fungi in other ionizing environments [28, 29].

Based on the empirical elemental formula for microbial (fungal) biomass [53] and the water content thereof [50], the theoretical MAC of *C. sphaerospermum* biomass at 150 MeV (the average cumulative energy of the Martian radiation environment, cf. supplementary information 1, section E) [54] was determined (cf. supplementary information 3 for derivation). This allowed µ_*Fungus*_ and the shielding potential of melanized biomass to be compared to other (theoretical) attenuation capacities of common aerospace construction materials and those considered for passive shielding against Space radiation (cf. supplementary information 1, section E). Both forms of the pigment common in melanized fungi (eumelanin and DHN-melanin) are comparatively effective radiation attenuators (µ_Melanin_ = 0.046 cm^-1^ at 150 MeV). Consequently, the predicted high attenuation capacity of melanized biomass at 150 MeV (µ_Fungus_ = 0.023 cm^-1^, compared to non-melanized fungal biomass with µ_Biomass_ = 0.016 cm^-1^) is a result of the high LAC of melanin.

Materials with large LACs typically have high densities and are therefore heavy (e.g., steel, cf. supplementary information 3). *C. sphaerospermum*, however, has comparatively large innate LAC and a (for organic compounds) low density, as does melanin. Due to the saprotrophic nature of the fungus, *C. sphaerospermum* has the ability to utilize virtually any carbon-based biomass for growth; on Mars this could be cyanobacterial lysate and/or organic waste, both of which have previously been proposed as feedstock for biotechnology [18, 55]. The radiotrophic nature of *C. sphaerospermum* is tantamount with radioresistance – it may thus be able to endure increased radiation exposure, effectively posing a self-regenerating radiation shield. Further testing, accompanied by dosimetry could substantiate this thesis and may also shine light on the impact of microgravity on the phenotype of these micro-fungi in a Space environment. It may also prove worthwhile to study other melanized fungi in a Space radiation context like e.g., *Wangiella dermatitidis* and *Cryptococcus neoformans*, which have been investigated for their radiotrophy and radioresistance in other Earth-based studies [28].

Various melanins have been investigated for their ability to protect against radiation, ionizing and non-ionizing, and found to be highly capable in that regard [23, 34, 48, 56-58]. Further, melanin-based composites can achieve synergistic improvements to radiation attenuating capacity. The MAC curve of a melanin-bismuth composite, for instance, is about double that of lead at energies in a low MeV range [59]. Another example of an enhanced, melanin-based radiation protection agent is selenomelanin: it was found that under increased radiation, nanoparticles of the compound could efficiently protect cells against cell cycle changes [60]. As a biological compound, natural melanin may be readily available through ISRU by means of biotechnology. In future, blends or layers of bio-derived melanin with other materials, analogous to the concept of Martian ‘biolith’ [61], may yield composites that more efficiently shield against radiation. This also opens the opportunity for melanin to be used as a constituent in fibrous composites for radiation shielding [62], to be used in textiles, such as EVA-suites or inflatable spacecraft and habitats. Advanced additive manufacturing technologies such as 3D-bioprinting, may ultimately even allow the creation of smart ‘living composite’ materials that are adaptive, self-healing and largely autonomous [63]. As live biomass has large water content, whole cells may thus present an excellent passive shield in particular for GCR. This will, however, require extensive further theoretical as well as experimental studies. Section G in supplementary information 1 contains additional remarks on this topic.

Regardless of how effective a radiation attenuator may be, passive shielding against GCR is ultimately always limited by mass [11, 64]. To increase density and thus the LAC, fungal biomass or melanin itself could be integrated with *in-situ* resources that are abundant at destination, such as regolith. In a case study we estimated that a ∼ 2.3 m layer of melanized fungal biomass (7.2% [w_melanin_/w_CWW_] melanin-content) would be needed to lower Martian radiation levels to those on Earth (from 234 mSv/a to 6.2 mSv/a [7, 8, 54]), whereas an equimolar composite of melanin and Martian regolith would only require a layer of ∼ 1 m for the same reduction of radiation. For comparison, in case of pure Martian regolith, about 1.3-times the thickness would be required to absorb the respective dose equivalent. When conducting the same analysis with purely non-melanized fungal biomass, a thickness of ∼ 3.5 m is required for the same shielding effect (reduction of radiation by 97%). Table 1 provides an overview of the inputs and outputs of this basic assessment, details can be found in supplementary information 1, section E.

**Table 1:** Comparison of radiation attenuating capacity of *in-situ* resources on Mars with melanized composites. Attenuation coefficients for the compounds were obtained from the [NIST XCOM](https://physics.nist.gov/PhysRefData/Xcom/html/xcom1.html) database [42], with buildup-factors generated by the RadPro Calculator [65], based on molecular formulas and/or densities for the respective materials as referenced, unless trivial or noted otherwise.

| Material and literature source for molecular formula and density | Mass Attenuation Coefficient '$\mu/\rho$' [$\text{cm}^2/\text{g}$] at 150 MeV * | Linear Attenuation Coefficient '$\mu$' [$\text{cm}^{-1}$] at relative material density | Required thickness [cm] to reduce Martian radiation levels by $\sim 97\%$ § |
| --- | --- | --- | --- |
| Water | 0.0149 | 0.0149 | 367 |
| Martian regolith [66] | 0.0268 | 0.0407 | 128 |
| Martian regolith + melanin (equimolar mixture) [67] | 0.0288 | 0.0518 | 106 |
| Non-melanized fungus<sup>§</sup> [53] | 0.0141 | 0.0155 | 351 |
| Melanized fungus<sup>#</sup> | 0.0213 | 0.0234 | 232 |

\* cumulative radiation environment on the surface of Mars [54]; § from 234 mSv/a, the average radiation dose on Mars [54], to 6.2 mSv/a the average radiation dose on Earth [7, 8]; § based on an empirical elemental formula for the biomass of baker's yeast [53], adjusted for 60% water content [50]; # based on 92.8% [w/w] non-melanized fungal biomass (as in §) and 7.2% [w/w] melanin content (DHN-melanin).

## Conclusion

With a basic experimental setup implemented as a small single payload, it could be shown that the dematiaceous fungus *C. sphaerospermum* can be cultivated in LEO, while subject to the unique microgravity and radiation environment of the ISS. Growth characteristics indicated an advantage of the on-orbit experiment as compared to the ground-control, potentially associated with beneficial effects of Space radiation, as has been suggested in analogous Earth-based studies. Further, monitoring of the radiation levels indicated that the melanized fungal biomass may have a radiation shielding effect. Attenuation of radiation was consistent over the whole course of the 30-day experiment, allowing a scenario-specific linear attenuation coefficient for *C. sphaerospermum* to be determined. This was further used to approximate the melanin content of the biomass, which corresponded well with literature. Based on the melanin content, the theoretical radiation attenuation capacity of fungal biomass could be put into perspective at constant equivalent ionizing radiation energy levels: melanized biomass and melanin containing composites were ranked effective radiation attenuators, emphasizing the potential melanin may holds as component of passive radiation shields.

Being living organisms, melanized micro-fungi self-replicate from microscopic amounts, which opens the door for ISRU through biotechnology and may allow significant savings in up-mass. Often nature has already developed surprisingly effective solutions to engineering and design problems faced as humankind evolves-biotechnology could thus prove to be an invaluable asset for life-support and protection of explorers on future missions to the Moon, Mars and beyond.

## Abbreviations

B: buildup-factor
CDW: cell dry-weight
CPM: counts per minute
CWW: cell wet-weight
DHN: 1,8-dihydroxynaphthalene
GCR: galactic cosmic radiation
HZE: high atomic number and energy
ISRU: *in-situ* resource utilization
ISS: international space station
LAC: linear attenuation coefficient
LEO: low earth-orbit
MAC: mass attenuation coefficient
PDA: potato dextrose agar
RT: room temperature.

## Acknowledgments

As members of ‘Team Orion’ Xavier R. Gomez, Srikar P. Kaligotla, Finn H. Poulin, and Jamison R. Fuller were instrumental for the conception and planning of the experiment. We extend our thanks to the Higher Orbits Foundation for providing funding for this project through the ‘Go For Launch!’ program, and to Space Tango, for the technical solution, logistics, and implementation of the experiment. Specifically, our gratitude goes out to Michelle Lucas of Higher Orbits and Gentry Barnett of Space Tango. Additional acknowledgements go to Kerry Lee of the NASA SRAG for help with obtaining dosimetric data for the timeframe of the experiment.

## Author contributions

GKS and XRG, and colleagues of ‘Team Orion’ conceived of the idea for the study in 2018 and composed the proposal for funding. Based on initial results, GKS compiled a draft-report with support from XRG, that served as basis for this paper. NJHA joined the team in early 2020, comprehensively analyzed the existing data, conducted supporting experiments, researched additional data, and wrote the manuscript with support from GKS. CK amended the study with regression models and statistical robustness analyses. All authors have read and approved the final version of the manuscript. The authors declare no competing interests. Correspondence and requests for materials should be addressed to NJHA and/or GKS.

## Footnotes

1 The melanin content of 7.2% [w_melanin_/w_CWW_] for wet biomass of *C. sphaerospermum* is approx. equivalent to a content of 21.5% [w_melanin_/w_CDW_] for dry biomass (based on a water content of 60% [49]). This appears to be in conformity with reported values for melanin contents of other microscopic fungi, which are in the range of 11.2% and 31.5% [w_melanin_/w_CDW_] [50, 51].

## Notes

### Competing Interest Statement

The authors have declared no competing interest.

### Summary of Updates

This revision of the manuscript details more thorough statistical and theoretical analyses, with revised numbers to support a similar conclusion, along with a corrected author list.

