## Supplemental Material 1 for "Growth of the Radiotrophic Fungus *Cladosporium sphaerospermum* aboard the International Space Station and Effects of Ionizing Radiation"

**Supplementary to: "A Self-Replicating Radiation-Shield for Human Deep-Space Exploration: Radiotrophic Fungi can Attenuate Ionizing Radiation aboard the International Space Station"**

### A. Microorganism Species

The dematiaceous (closely related to 'black yeast') fungus *Cladosporium sphaerospermum* was originally discovered in 1886 by Albert Julius Otto Penzig [1] and is known to be saprotroph, xerotolerant and halotolerant; its radiotrophy (ability to utilize ionizing radiation for metabolic functions), however, was first discovered following the disaster at the Chernobyl Nuclear Power Plant in 1986, when the melanized fungi thriving in the surrounding highly radioactive environment were investigated [2]: fungal samples taken from Chernobyl that were exposed to radiation levels approximately 500-times higher than background level grew significantly faster than those not exposed to radiation [3]. These organisms continued to grow towards the source of radiation, another indication for the affinity of the organism towards ionizing radiation and thus radiotrophy [4]. Radiosynthesis is perceived analogous to photosynthesis, except that where the intricacies of photosynthesis have been extensively studied and are generally accepted, the precise mechanism of radiosynthesis remains unsettled. Nevertheless, it has been suggested that ionizing radiation alters the chemical properties of melanin in the cell, which leads to increased rates of electron-transfer, allowing for a net energy-gain to be used to reduce carbon [3, 5] (analogous to how the chlorophyll in photosynthesis captures the energy of light to be stored chemically). Unlike melanocytes in humans, in fungi the melanin is crystallized on the cell-walls [6]. This provides protection of the cells from oxidizing agents (free radicals) generated by different forms of ionizing and non-ionizing radiation (UV-light), giving the mold an ecological advantage in extreme environments on Earth, natural as well as non-natural. The expectation is that these capabilities may also prove advantageous in a Space environment. Therefore, loosely analogous to an existing Earth-based study [4], *C. sphaerospermum* was selected as subject for an experiment on the International Space Station (ISS).

### B. Methodology and Experiment

<u>Absorption Spectrum of Radiation Sensors</u>

Ionizing cosmic radiation, or in other words radiation with enough kinetic or electromagnetic energy to strip electrons from an atom, comes in two forms (excluding photons from the classification): wave radiation (electromagnetic waves) and particle radiation, like e.g. $\alpha$- and $\beta$-particles or HZE ions [7]. In the electromagnetic spectrum, ionizing (wave-) radiation is characterized by energies ranging from a few hundred eV to about 1 MeV, but can also be much higher for cosmic radiation. The PocketGeiger Type5 is designed to measure wave radiation, specifically of X- and $\gamma$-type [8]: at 23°C the PIN photodiode of the X100-7 SMD (First Sensor AG, Berlin, Germany) is sensitive for $\gamma$-radiation with energies of 3-20 keV, with highest absorbance between 5.5 keV and 10 keV [9].

<u>Levels and Energy of Measured Radiation</u>

Based on the total cumulative recorded radiation counts over the 622 hours of the experiment, an average of 149 CPM (counts-per-minute) was determined. With reported radiation levels for the ISS of $\approx$ 144 mSv/a ($\triangleq$ 0.274 µSv/min) or 58 mGy/a ($\triangleq$ 0.11 µGy/min) [10], respectively, a single count thus corresponds to an average of approx. 5.6 nSv or 2.2 nGy, respectively.

Additionally, daily dosage data obtained from the Destiny module (US Lab) of the ISS for the time of the experiment was correlated with the radiation counts from the experiment (cf. supplementary information 2). The trend followed most closely the fraction of the radiation attributed to the South Atlantic Anomaly (SAA), while no direct correlation was observed between the radiation counts and dosage attributed to Galactic Cosmic Radiation (GCR). This indicates that electrons in the range of hundreds of keV and energetic protons with energies exceeding 100 MeV, as are typical for the inner Van Allen belt, where likely the primary forms of energetic particles picked up by the employed radiation sensors, in addition to secondary radiation in the absorbance range of the radiation sensors.

<u>Flight Hardware and Experimental Implementation</u>

The flight-hardware was packaged as a 2U (double standard-size) CubeLab™ (sealed) module (4"×4"×8"), which has a volume of 103.4 in$^3$, is air-tight, and provides up to 20 W of power. The assembled unit is shown in figure S1.

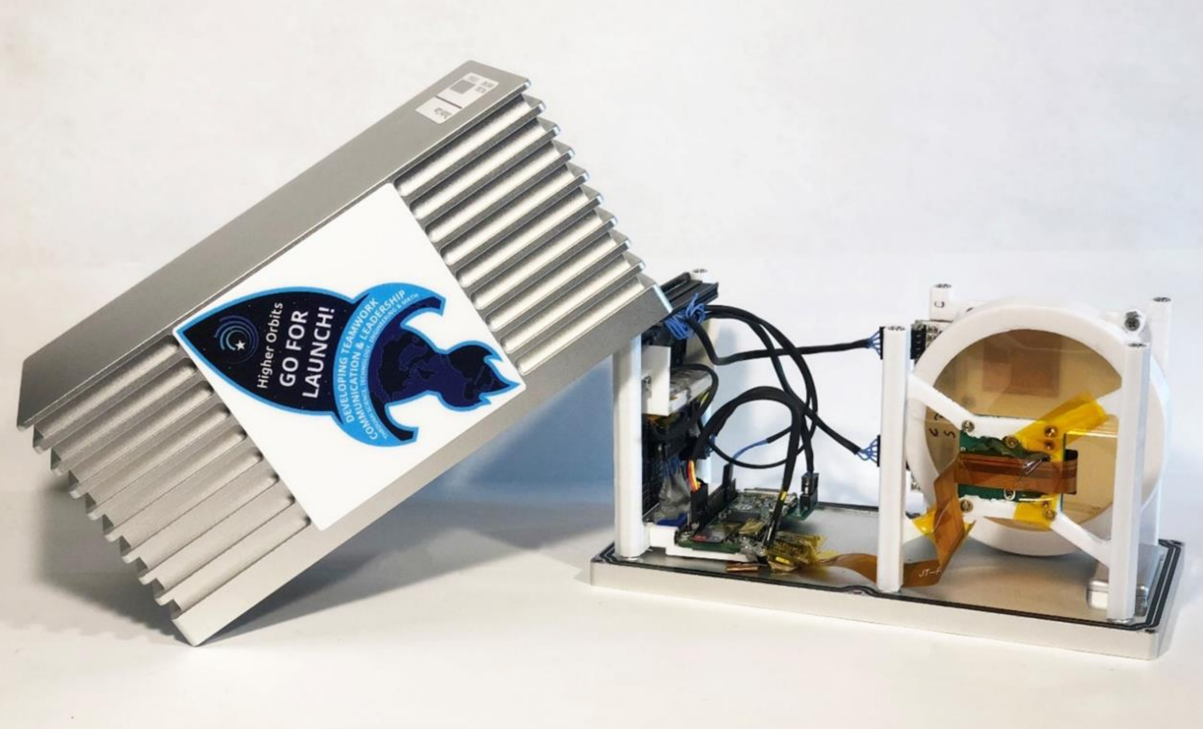


**Figure S1***:* Fabricated flight hardware unit, packaged in a 2U Space Tango CubeLab™. Media Courtesy: Space Tango, Inc.

The assembled flight-hardware was calibrated and vetted before flight; in particular, consistency of the two radiation sensors was confirmed, so that no deviation existed between them. Furthermore, it was ensured that no fungal growth would occur prior to activation of the hardware ($t_0$) by means of cold-stow while enroute to the ISS. Preliminary experiments confirmed that nominal growth occurred over several days of cold storage of inoculated medium. On-orbit the unit was oriented in a way that the Petri dish faced away from Earth to maximize radiation exposure.

### C. Evaluation of Microbial Growth

Growth of *C. sphaerospermum* was characterized based on the relative brightness of the time-series photos of the fungus grown on solid medium. MATLAB was utilized to automate evaluation of the images by means of "HSV colormap array". For this purpose, a representative section of the fungus-covered agar with high contrast was chosen, as indicated in figure S2 I for the on-orbit experiment. The obtained values were standardized to an optical density (OD) of 0 at $t_0$ and normalized to the average brightness value of the last 4-8 hours of the experiment to a maximum OD of 1. Plotted over time, relative growth-curves were thus obtained (cf. supplementary information 2).

The data for the ground-control experiments were processed equivalently, integrating multiple replicates. Specifically, for the ground-control, three individual experiments were carried out. Each experiment contained three replicates (separate Petri dishes), the OD-data of which was combined to obtain robust growth-curves and the deviation for each experiment (cf. supplementary information 2).

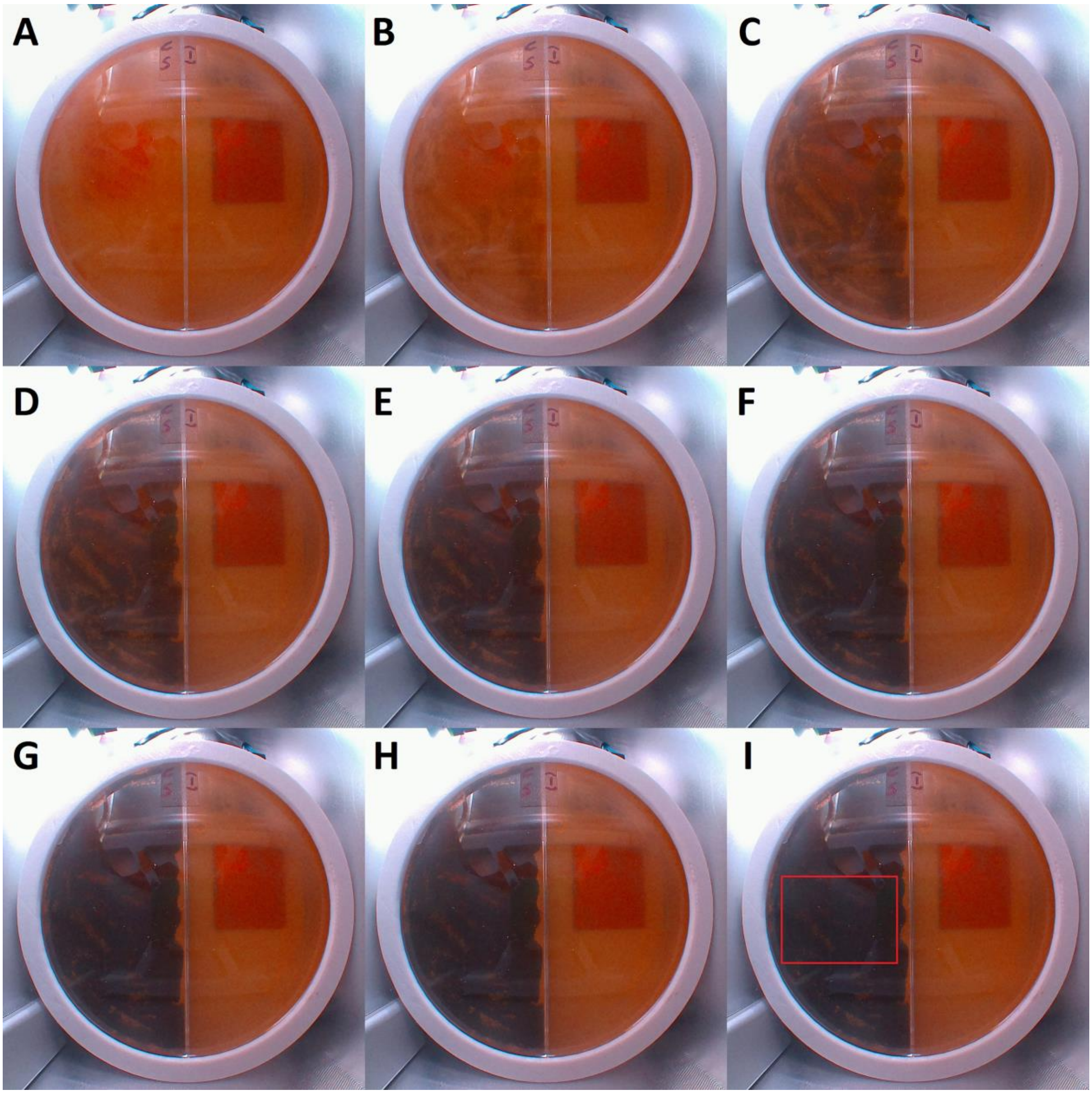


**Figure S2: Photographic data of fungal growth**. Images A - I of the on-orbit experiment show fungal growth on the agar of one side of the split Petri dish in intervals of 6 hours starting with A: $t_0 = 0$ hours continuing until I: t = 48 hours. The red frame in the last picture (I) indicates the section of the picture used to derive the growth data by means of picture brightness, exemplary for all pictures.

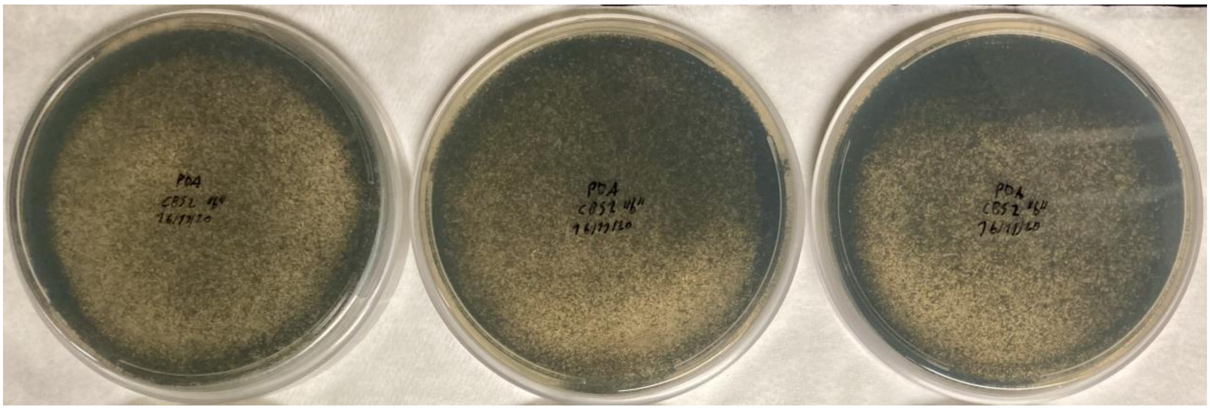

**Figure S3: Photo of ground-control experiment**, showing growth at 30°C 14 days after inoculation.

### D. Linear Attenuation Analysis

Definition and Background of Calculations

A linear attenuation coefficient (LAC, symbol "$\mu$") is a constant, given in per unit thickness of a material. It describes the fraction of attenuated incident uncharged particles in a monoenergetic beam [11]. Therefore, it can be used to determine the probability that ionizing (wave or particle) radiation will be scattered by a material, scaled on a linear attenuation index [12]. An LAC is a material-intrinsic property and can be used to assess the ability of a material to block radiation, as change in energy levels from initial to final radiation dose over the length of the barrier (a form of Lambert's law). A mass attenuation coefficient (MAC, symbol $\mu_m$) represents the single-beam attenuation capacity of a volume to absorb wavelength radiation at any density.

The interactions of charged particles such as protons and HZE ions with matter are complex and are best described by their stopping power. For attenuation coefficients to be extended to charged particles, and not ordinarily ignore secondary radiation, it must be accounted for, which can be done by means of buildup-factors [13]. These accounts for interactions associated with photons such as the Compton and photoelectric effects [14], but not Bremsstrahlung and Spallation, nor is the excitation or emission of photoelectrons in a mass respected. Buildup-factors (symbol "B") are specific for each material and electron volt energy and are generated based on relative energy absorption. They are derived from the number of mean free paths (relaxation length, symbol "R") for a system, by accounting for the energy level of relativistic particles. Therefore, calculations based on linear attenuation that respect B-factors are valid for a monoenergetic radiation environment [15],[1] where all photonic secondary radiation is accounted for but secondary radiations associated with charged particles such as Bremsstrahlung, and Spallation are ignored and do not take into account the energy density of spalled neutrons caused by indictment GCR protons. Such spalled particles are produced within and emanate from spallation zones adjacent to the outer free surface of a body [16]. Moreover, the energy density of such spalled particles on the Surface of Mars at e.g., 150 MeV is less than approximately 1500 neutrons per second per gram of material [17]. While the final radiation doses may be underestimated, it should be noted that the associated error is low, as the majority of galactic charged particles are not energized enough to create photoelectrons nor additional secondary radiation within or adjacent to a mass [18].

The calculations that supported the present analysis were based on following equations:

Eq. 1 $\quad I = B \times I_0 \times e^{-\mu \times x}$

Eq. 2 $\quad \mu = \sum w_i \times \mu_i$

Eq. 3 $\quad \mu_m = \mu / \rho_m$

where:

- $I_0$ = initial radiation intensity
- I = transmitted radiation intensity
- B = buildup-factor (unitless)
- $\mu$ = linear attenuation coefficient in $cm^{-1}$
- ρ = density of material in g/cm
- $x$ = thickness of material in cm
- $w_i$ = weight fraction of species $i$ (unitless)

[1] for multiple photon energies the correct number of scattered photons may be estimated for each energy individually as the sum of the weighted contributions

In Eq. 1 'I' represents the intensity of a beam at distance $x$ in dependence of the attenuation coefficient '$\mu$', the initial intensity '$I_0$' and, if applicable, a buildup-factor 'B' > 1. As an observation of the Beer-Lambert Law, Eq. 1 only holds true under certain assumptions [19].[2] Eq. 2 defines the working linear attenuation coefficient '$\mu$' of a mixture of different chemical species, where $w_i$ is the weight fraction of a species $i$. Eq. 3 describes the relationship between linear attenuation coefficients, $\mu$, and mass attenuation coefficients '$\mu_m$'. Since linear attenuation coefficients are dependent on density, mass attenuation coefficients are commonly reported [20]; LACs can be derived if the relative mass density '$\rho_m$' of a given material is known [21].

### Buildup-factors

There exists a quadratic dependence between linear/exponential attenuation coefficients and buildup-factors across a wide range of materials, with independent relationships for the number of mean free paths, R [13]. Accordingly, B-factors were interpolated, based on data from the RadPro Calculator [22] for the mean photon energy around the International Space Station (R = 0.5 cm), and for the Martian radiation environment (R = 3.629 cm). This allowed B to be derived for any linear attenuation coefficient at the specific radiation environment of interest, as per figure S4 A & B.

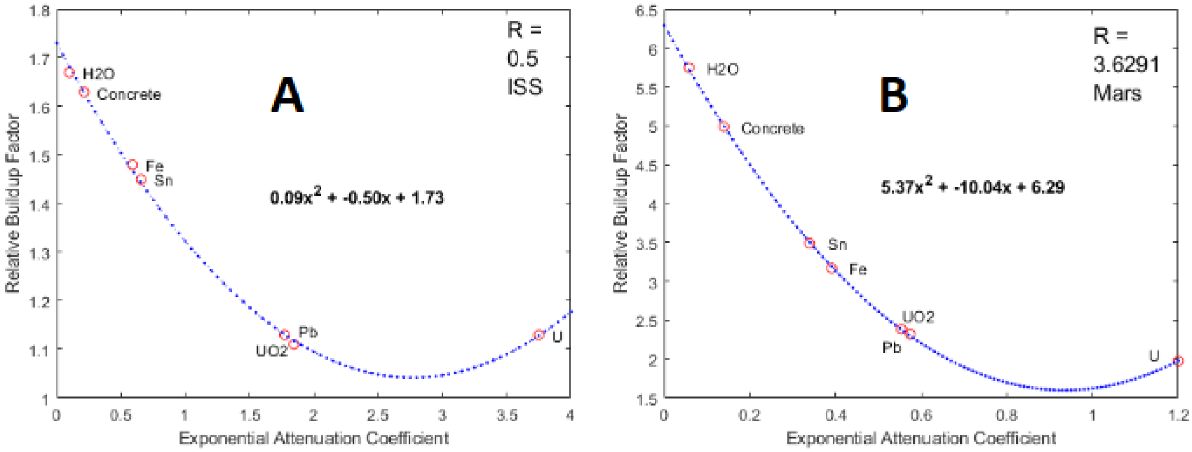


**Figure S4: Quadratic interpolation of buildup-factors for the radiation environment of the ISS (A) and Mars (B).** R, in centimeters, was correlated to the unique radiation environments surrounding Mars (150 MeV) [17] and the ISS (50 MeV) [23], as derived from a table of LACs, B-factors, and R-values from literature [13]. Fe = iron, $H_2O$ = water, Pb = lead, U = Uranium, $UO_2$ = Uranium dioxide, Sn = Tin.

[2] of the six assumptions, the most relevant are that electromagnetic coupling must be excluded and radiation must pass through a homogenous attenuating medium

Attenuation Analysis

The radiation attenuation, as per the relative difference between the slopes of the linear regression of cumulative radiation counts from the experiment and negative-control, was used to infer a ratio between initial and transmitted radiation intensity such that $I_{Control} / I_{Experiment} = 1.00871$, where $I_{Control}$ is representative of $I_0$ (with respective absolute ratios of the initial and final phases of 1.01552 and 1.02423 the relative difference is 1.00871). Since the fungal layer shielded only one side of the sensors from radiation the ratio can be doubled such that $I_0 = 1.01742 \times I$, and Eq. 1 becomes:

$$I = B \times (1.01742 \times I) \times e^{-\mu \times x} \Leftrightarrow (B \times 1.01742)^{-1} = e^{-\mu \times x}$$

With the approximated thickness of the fungal lawn (≈ 0.167 cm), and B-factors for 5.5 and 10 keV ($B_{5.5} = 3.84$ and $B_{10} = 1.43$), respectively,[3] the MAC of *C. sphaerospermum*, $\mu_m$, was determined accordingly:

for 5.5 keV: $(3.84 \times 1.01742)^{-1} = e^{-\mu \times 0.167\ cm} \Leftrightarrow \mu_{Fungus5.5} = 8.16\ cm^{-1}$

for 10 keV: $(1.43 \times 1.01742)^{-1} = e^{-\mu \times 0.167\ cm} \Leftrightarrow \mu_{Fungus10} = 2.245\ cm^{-1}$

At lower energies, LACs do not only grow exponentially larger, but small changes in energy yield large changes of the value, which needs to be taken into account in any further analyses. Hence, $\mu_{Fungus}$ is reported with a standard deviation of the two boundary values as $5.2 \pm 4.2\ cm^{-1}$.

Beyond being a measure of this fungus' ability to shield against ionizing radiation, the LAC of *C. sphaerospermum* can be used to estimate its melanin content, based on Eq. 2:

$$\mu_{Fungus} = \frac{m}{1} \times \mu_{Melanin} + \frac{1-m}{1} \times \mu_{Biomass}$$

where:

- $m$ = mass of melanin as fraction of any given amount of biomass.
- $\mu$ = LACs for melanin and fungal biomass, retrieved from NIST-XCOM.

This requires the approximation that fungal biomass can be defined as a homogeneous aqueous solution, with a single molecular sum-formula: the elemental composition of non-melanized wet biomass was determined as per supplementary information 3, based on the empirical formula for dry biomass [24] and average water content [25] of fungi. Evaluated with the boundary LACs at 5.5 keV and 10 keV, the melanin content of wet biomass was thus estimated as follows:

for 5.5 keV: $2 \times 8.16\ cm^{-1} = \frac{m}{1} \times 31.32\ cm^{-1} + \frac{1-m}{1} \times 15.15\ cm^{-1} \Leftrightarrow m = 0.072356$

for 10 keV: $2 \times 2.245\ cm^{-1} = \frac{m}{1} \times 8.99\ cm^{-1} + \frac{1-m}{1} \times 4.14\ cm^{-1} \Leftrightarrow m = 0.072165$

The melanin content of *C. sphaerospermum* under the given cultivation conditions aboard the ISS therefore seems to be around 7.2% [w/w]. This is realistic according to the following feasibility assessment: the fraction of (dry) cell-mass that is cell-wall is around 25% [w/w] [24, 26]. Hence, if half (50% [w/w]) of the (dry) cell-wall was melanin, the maximum fraction of (dry) cell-mass that is melanin would be around 12% [w/w] or, when factoring in a water content of ~ 60% [w/w] [25], around 7% [w/w] for wet cell-mass. Similarly, the approx. melanin content of dried fungal biomass can be determined as ~ 21.5% [w/w].

---

[3] separate Buildup-factors were determined for the boundary energies of the optimal absorption range of the PocketGeiger Type5 (cf. section B)

### E. Contextual Attenuation Comparison

The equivalent dose that incorporates all aspects of Martian ionizing radiation, as described by a quality-factor of 3.05, is 233.6±36.5 mSv/a [17]. GCR contributes approximately 122 to 153 MeV to the Martian radiation environment, while SEP events, on average, account for ~ 11 MeV. Photons in the Martian atmosphere contribute another 1.2 to 1.7 MeV. Hence, the combined total energy of the Martian radiation environment ranges from 134 to 166 MeV so that the average cumulative radiation intensity on the surface of Mars is approx. 150 MeV [17]. This generalization is acceptable in the present context since the LAC curves at the energies of interest are relatively flat and the mathematically equivalent energy value of an approximately homogeneous radiation environment accounts for mass, momentum, and electromagnetic energies, from which LACs and B-factors can be derived.

The integration of microbial melanin with compounds by chelating certain metals can effectively increase the overall attenuation capability of the composite material in high-stress γ-ray and GCR environments [27]. However, for ISRU, it is most meaningful to assess the attenuation capability of a melanin composite when combined with a resource that is abundant at destination, such as regolith. Let material X represent an equimolar mixture of melanin and Martian regolith, such that its linear attenuation coefficient, $\mu_X$, at an energy of 150 MeV and a medial density of melanin and regolith [28] is explicitly defined. With an average radiation dose on Earth of $I_{Earth}$ = 6.2 mSv/a [29] and $I_{Mars}$ = 233.6 mSv/a, Eq. 1 can be rewritten to expresses the ability of material X to lower the equivalent dose of Martian radiation to Earth levels:

$$x = \ln[(I_{Earth} / I_{Mars})/B]/(-\mu_X)$$

In this scenario, the required material thickness $x$ of material X to lower Martian radiation levels by ~ 97% (from 233.6 mSv/a to 6.2 mSv/a) is approx. 1 m, depending on the range of energies on the Martian surface. To put the result into perspective, Table S1 was compiled, which contains the results of analogous calculations for various other materials common in aerospace or those considered as *in-situ* resources for radiation shielding on Mars. See supplementary information 3 for details and the calculations of composition of mixtures.

**Supplementary to: "A Self-Replicating Radiation-Shield for Human Deep-Space Exploration: Radiotrophic Fungi can Attenuate Ionizing Radiation aboard the International Space Station"**

**Table S1: Comparison of radiation attenuating capacity of different materials common in spaceflight or proposed for radiation shielding of spacecraft and surface-habitats** with melanized/non-melanized biomass in a reference-scenario simulating deep Space radiation conditions. Attenuation coefficients were generated from the NIST XCOM database [14], based on molecular formulas and/or densities for the respective materials as referenced, unless trivial or noted otherwise.

| **Material and literature source for molecular formula and density** | **Mass Attenuation Coefficient '$\mu/\rho$' [$cm^2/g$] at 150 MeV *** | **Linear Attenuation Coefficient '$\mu$' [$cm^{-1}$] at relative material density** | **Required thickness [cm] to reduce Martian radiation levels by ~ 97% $** |
|---|---|---|---|
| Water | 0.0149 | 0.0149 | 355 - 376 |
| Non-melanized fungus§ | 0.0141 | 0.0155 | 340 - 360 |
| Melanized fungus# | 0.0213 | 0.0234 | 225 - 238 |
| Eumelanin | 0.0307 | 0.0463 | 113 - 120 |
| DHN-melanin | 0.0308 | 0.0464 | 113 - 119 |
| Lunar regolith [30] | 0.0229 | 0.0344 | 153 - 162 |
| Martian regolith [31] | 0.0278 | 0.0423 | 124 - 131 |
| Melanized Martian regolith † | 0.0317 | 0.0571 | 102 - 108 |
| Aluminosilicate [32] | 0.0241 | 0.0554 | 94 - 100 |
| Aluminum | 0.0264 | 0.0713 | 73 - 77 |
| Stainless Steel (301) | 0.0450 | 0.3690 | ~ 13 |
| HDPE | 0.0145 | 0.0141 | 374 - 397 |
| PET | 0.0161 | 0.0222 | 237 - 251 |
| Kevlar [33] | 0.0159 | 0.0229 | 230 - 244 |
| Carbon Fiber Cloth | 0.0150 | 0.0290 | 181 - 192 |
| Beta Cloth [34] | 0.0184 | 0.0403 | 130 - 138 |

* cumulative radiation environment on the surface of Mars [17]; $ from 233.6 mSv/a to 6.2 mSv/a, the average radiation doses on Mars [17] and Earth [29], respectively; § based on an empirical elemental formula for the biomass of baker's yeast [26]; # based on 92.8% [w/w] non-melanized fungal biomass (baker's yeast, adjusted for 60% water content [26]), and 7.2% [w/w] melanin content (DHN-melanin); † based on an equimolar mixture of Martian regolith and (DHN)-melanin. HDPE = high-density polyethylene; PET = polyethylene terephthalate.

### F. Data Confidence and Reliability of Results

For Geiger counters and dosimeters alike, the percent error of the measurements can change when the radiation levels change (cf. supplementary information 2, plot of radiation count vs. noise signal). Because of indeterminable absolute error for each individual read, an average percent error is assigned to instruments like the sensor used in this experiment. Since here not the absolute level of radiation, but the difference between the two sensors was determining, it was sufficient to establish consistency of the two devices prior to flight and in the initial phase of the experiment on-orbit. Further, with ~ 24k data points collected over the 622-hour runtime of the experiment and an $R^2$ of the linear regressions > 0.99, robustness of the results is warranted. Hence, the determined 1.68% of radiation attenuation seem to be reliable. Nevertheless, the assumption that radiation intensity on the ISS is not dependent on a vector may involve an error, as the ISS does not "spin" while orbiting, so that the "bottom" facing Earth may receive a different radiation dose than the "top". Therefore, the true attenuation capacity may be somewhere between 0.84% and 1.68%.

Buildup-factors are influential for highly energetic radiation and account for overestimations in linear attenuation coefficients. They are therefore used in calculations relying on the Beer-Lambert law. B-factors are able to return the calculation to within 2% accuracy of the actual radiation environment [13], which is sufficient for the qualitative statements of this study. Subject to the described parameters and assumptions, the presented calculations therefore allow a fair appraisal of the potential shielding capacities. Nevertheless, to holistically describe the real (off-)world radiation environment, due to the complex nature of GCR and vicissitude of space weather, and to precisely study the shielding properties of melanin containing materials, tools like OLTARIS [35] and SPENVIS [36] and/or Monte Carlo simulations [37] (using platforms like e.g. GEANT4 and CREME96) [38] are sensible to model and further validate findings. This is, however, out of the scope of this study. Nevertheless, we feel it is important to mention such analyses which go hand-in-hand with experimental studies, like Galactic Cosmic Ray Simulation (GCRSim) [39] at the NASA Space Radiation Laboratory, and a deeper understanding of the health risk that space radiation poses for human crews.

### G. Expanded Discussion on ISRU

One possible application of *in-situ* resource utilization (ISRU) on Mars is to leverage existing resources to increase availability of habitation, in order to significantly reduce the required haul of construction material and/or prefabricated structures from Earth as well as to break the supply chain and provide independence and redundancy. For this, a multitude of different approaches exists, many of them are still inhibited by the extent of initially required critical infrastructure [40, 41]. More recently, autonomous 3D-printing of infrastructure relying on composites with Martian regolith has been proposed [42], however, this also still requires significant up-mass of auxiliary equipment to allow for e.g., stripping and processing of Martian topsoil, as well as the raw material for the binding resin. However, if the binding material could also be derived or produced on-site, an additive manufacturing method may become immediately more feasible.

While bio-based methods are still in early stages of development [43], fungal structures have already been targeted by NASA through investigation of mycotecture as a potential technology for production of structural components [44]. Further expanding the range of fungi being tested for growth on regolith simulants would yield information on the feasibility of *C. sphaerospermum* composites with ISRU obtained resources to bolster radiation shielding capacity of habitation. Ultimately "Engineered Living Materials" (ELMs) [45] may provide the game-changing solution for the most pressing issues of deep-space exploration (up-mass and radiation). This would allow the tailored construction of structural and supporting components, possibly in combination with 3D-bioprinting, which has already been shown to be feasible, also with fungal mycelium [46].

A passive shield that is an effective attenuator as well as available and readily deployable at destination Mars would significantly contribute to a solution for radiation shielding of habitats, when looking to establish a permanent foothold on the fourth rock from the sun. In future, manufacturing of "melanin-infused" ceramics and alloys into spacecraft, may increase radiation protection capability while keeping the wall-thickness and weight of the vessel relatively constant: instead of increasing the strength of construction materials such as steel or aluminum, which would result in substantial increase of mass (~ 7 g/cm$^3$ for standard steel at Earth gravity), materials could be "reinforced" with melanin, e.g. woven in with melanin as with fabrics, thus maintaining a low area density [47]. The possibility to polymerize melanin itself [48] may even lead to the development of advanced high-performance (plastic) materials based on these natural pigments.
